# BayesSpace enables the robust characterization of spatial gene expression architecture in tissue sections at increased resolution

**DOI:** 10.1101/2020.09.04.283812

**Authors:** Edward Zhao, Matthew R. Stone, Xing Ren, Thomas Pulliam, Paul Nghiem, Jason H. Bielas, Raphael Gottardo

## Abstract

Recently developed spatial gene expression technologies such as the Spatial Transcriptomics and Visium platforms allow for comprehensive measurement of transcriptomic profiles while retaining spatial context. However, existing methods for analyzing spatial gene expression data often do not efficiently leverage the spatial information and fail to address the limited resolution of the technology. Here, we introduce BayesSpace, a fully Bayesian statistical method for clustering analysis and resolution enhancement of spatial transcriptomics data that seamlessly integrates into current transcriptomics analysis workflows. We show that BayesSpace improves the identification of transcriptionally distinct tissues from spatial transcriptomics samples of the brain, of melanoma, and of squamous cell carcinoma. In particular, BayesSpace’s improved resolution allows the identification of tissue structure that is not detectable at the original resolution and thus not recovered by other methods. Using an *in silico* dataset constructed from scRNA-seq, we demonstrate that BayesSpace can spatially resolve expression patterns to near single-cell resolution without the need for external single-cell sequencing data. In all, our results illustrate the utility BayesSpace has in facilitating the discovery of biological insights from a variety of spatial transcriptomics datasets.

## Introduction

Single-cell RNA sequencing (scRNA-seq) achieves high-throughput and high-resolution profiling of gene expression, but traditional sample preparation results in the loss of spatial information. Knowledge of the spatial location of transcript expression can provide vital insights into biological function and pathology. Fortunately, technological advances have allowed for high-throughput profiling of gene expression while retaining spatial information^1^. Such spatially resolved transcriptomic profiling allows analyses to be made within the context of the biological tissue. Studies done on the Spatial Transcriptomics (ST) platform and the closely related Visium platform have already generated insights into diverse areas such as tumor heterogeneity^2,3^, brain function^4^, and the pathophysiology of sepsis^5^. The primary technological limitation of these spatial gene expression platforms is resolution, with the unit of observation being spots that are 100μm in diameter on the ST platform and 55μm in diameter on the Visium platform. As such, the number of cells within a spot may range from 1 to 200 depending on the biological tissue and the technological platform^6^. Alternative approaches include fluorescence *in situ* hybridization technologies, such as seqFISH and MERFISH, and other recently developed spatial sequencing methods, such as Slide-seq and ZipSeq^7–10^. While these methods provide increased resolution, most are lower throughput, rely on custom protocols, or are not widely available.

Presently, there is a need for new statistical methods for the analysis of spatial gene expression data that efficiently leverage the available spatial information. Clustering is an important step in the analysis of such data that allows downstream analyses such as cell type or tissue annotation and differential expression to provide unbiased biological insights. Existing analyses of spatial gene expression data often rely primarily on clustering methods for non-spatial RNA-seq data^2,4^. The additional spatial information available from ST and Visium can help address the challenges to analysis brought by sparsity and noise by smoothing over adjacent spots, which are more likely to have similar transcriptomic profiles. Zhu et al. (2018) proposed a hidden Markov random field model (HMRF) for clustering of low-resolution *in situ* hybridization data into distinct spatial domains by jointly modeling gene expression and the spatial neighborhood structure^11^. This approach was later adapted for use with high-throughput spatial transcriptomics data through the selection of spatially differentially expressed genes prior to clustering^12^. Another recently developed spatial clustering algorithm is stLearn, which uses deep learning features extracted from the histopathological images as well as the expression of neighboring spots to spatially smooth the data^13^.

Here, we propose BayesSpace, a fully Bayesian spatial clustering method that models a low-dimensional representation of the gene expression matrix and encourages neighboring spots to belong to the same cluster via a spatial prior (Fig. 1A). From a modeling perspective, BayesSpace allows for a more flexible specification of the clustering structure and error term than alternative approaches. From a user perspective, BayesSpace is accessible in that it takes the widely used Bioconductor SingleCellExperiment object as input^14^, does not require the additional task of preselection of marker genes, and involves minimal parameter tuning. Our method draws from previously developed spatial statistics methods for image analysis and microarray data^15,16^.

**Figure 1.**
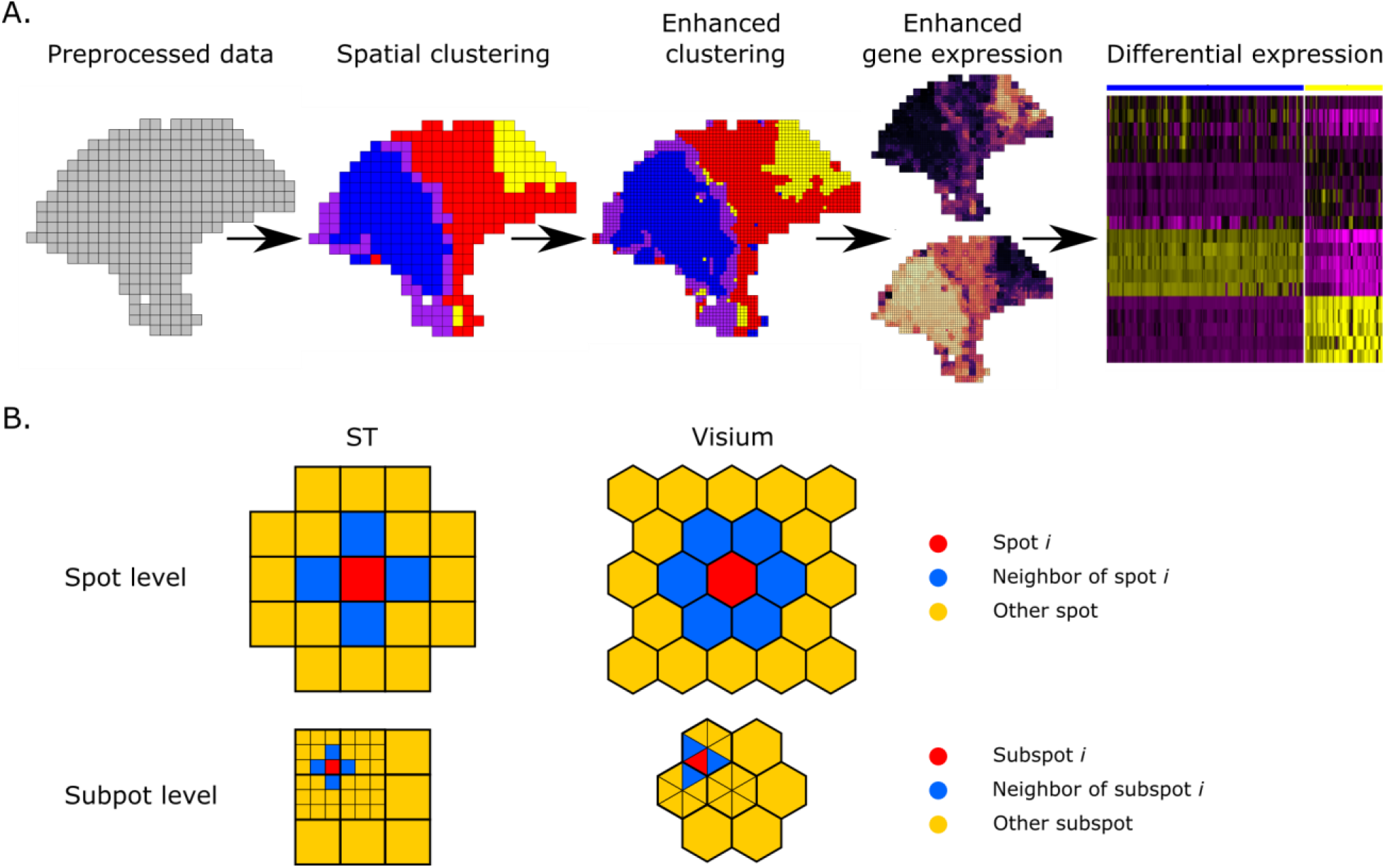
The BayesSpace workflow. (A) The BayesSpace workflow begins with preprocessed ST or Visium data. The data are spatially clustered to infer regions with similar expression profiles. These clusters can be refined via enhanced clustering to provide a higher resolution spatial map. Enhanced clustering also provides the basis for predicting gene expression at the higher resolution, which can be used in further differential expression analyses. (B) From geometric representations of the spatial distribution of spots in the ST and Visium technologies, neighbors can be identified for each spot based on shared edges (top). Each spot can be subdivided into subspots, which again have natural edge-based neighbors (bottom).

In addition, there is a need for spatial gene expression methods that address the relatively low resolution of the technology. Existing spatial gene expression deconvolution methods build upon bulk RNA-seq deconvolution methods and depend on additional scRNA-seq data^17–19^. While integration with scRNA-seq is appealing, it may be costly if using matched samples or introduce bias if using publicly available references. Furthermore, the deconvolved mixtures are still only spatially resolved at the original scale of the ST or Visium technology and the neighborhood structure of the cell types can’t be recovered. We extend BayesSpace to resolve expression at a subspot level by leveraging the spatial neighborhood structure (Fig. 1A). This enhanced resolution modeling – which approaches single-cell resolution with the Visium platform – does not require independent single-cell data and allows us to infer the spatial arrangement of the subspots.

Using several datasets we show that BayesSpace improves the identification of spatially distributed tissue domains through spatial clustering and enhances the resolution of gene expression maps. Furthermore, using *in silico* spatial transcriptomics datasets generated from aggregating single-cell RNA-seq, we show that BayesSpace can recover the true spatial structure at near single-cell resolution.

## Results

### Spatial clustering improves the identification of known layers in brain tissues

Recently, Maynard et al. (2020) presented Visium spatial expression profiles of twelve dorsolateral prefrontal cortex (DLPFC) samples, as well as manual annotations of the six cortical layers and white matter for each sample as part of the spatialLIBD package^4^ (Fig. 2A). Here, we evaluate BayesSpace’s ability to identify distinct layer-specific expression profiles and compare its performance to other spatial and non-spatial clustering methods. Specifically, we compare the performance of three non-spatial algorithms commonly applied to scRNA-seq data – *k*-means, mclust^20^, and Louvain^21^; two recently published spatial clustering algorithms – HMRF (as implemented in the Giotto package)^12^ and stLearn^13^; and the clustering partitions originally reported by Maynard et al. in the spatialLIBD package, which involve Walktrap clustering of spatial coordinates and PCs calculated from highly variable genes (HVGs) or known layer-specific marker genes. Following the methodology of Maynard et al., we use the adjusted Rand index (ARI) to quantify the similarity between cluster labels and the manual annotations, which are considered the ground truth.

**Figure 2.**
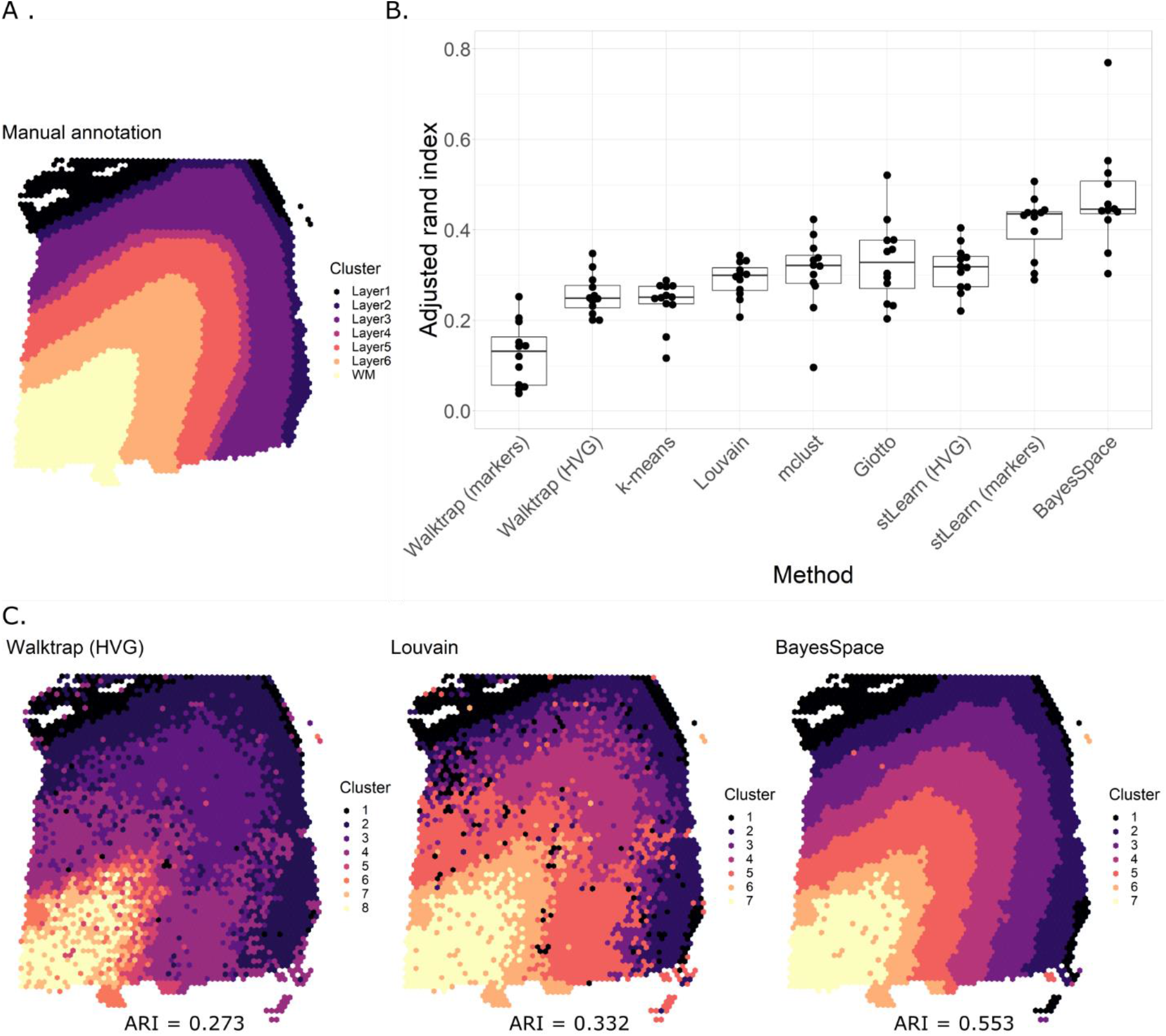
BayesSpace improves computational resolution of layers in the dorsolateral prefrontal cortex. (A) Ground truth. We highlight the manually annotated six DLPFC layers and white matter in sample 151673 from the spatialLIBD dataset. (B) Summary of clustering accuracy in all twelve samples. ARI is used to compare the similarity between cluster labels from each method against the manually annotated layers for all twelve samples. (C) Cluster assignments in sample 151673. The original cluster labels reported by Maynard et al. using Walktrap on HVGs (left), clustering labels from the non-spatial Louvain clustering algorithm (center), and BayesSpace spatial clustering labels (right).

BayesSpace substantially outperforms the original spatialLIBD clustering partitions, as well as all non-spatial clustering algorithms and spatial clustering methods developed for spatial transcriptomics data (Fig. 2B). BayesSpace and the non-spatial methods are applied on 15 PCs calculated from the top 2000 HVGs. The spatial clustering methods, Giotto and stLearn, are implemented based on the original authors’ recommended parameters (Supplementary Information). In sample 151673, we find that the Walktrap (HVG) clustering partition reported by Maynard et al. (ARI = 0.273) and the non-spatial Louvain clustering partition (ARI = 0.332) both exhibit substantial noise and lack of clear spatial separation between clusters (Fig. 2C). The white matter and inner layers do not cluster into distinct bands. In contrast, BayesSpace leverages spatial information to smooth the data and provides distinct layers of clusters (ARI = 0.553). The *t*-distributed error model that BayesSpace uses is particularly robust against outliers in the clusters, which may be caused by dropout events or other technical artifacts (Fig. S1).

### Increased resolution clustering leads to the identification of known tissue structures missed by other methods

We use BayesSpace to analyze a melanoma ST sample first annotated and described by Thrane et al^2^. Since the manual annotation identifies regions of melanoma, stroma, and lymphoid tissue and leaves an additional area unannotated (Fig. 3A), we run spatial clustering with four clusters (Fig. 3B). The resulting clusters correspond well with the manually annotated tissue types. Furthermore, the melanoma tissue is split into the central region of the tumor and an outer ring of mixed tumor and lymphoid tissue. BayesSpace enhanced spatial clustering provides a higher resolution map of the tissue types (Fig. 3C). Notably, the enhancement identifies lymphoid regions along the tumor border and possible immune infiltration into the tumor that cannot be discerned at the original resolution. These regions are also largely not identified by other clustering methods (Fig. S2A). While most clustering methods identified heterogeneity between the periphery and the center of the tumor, only Giotto and BayesSpace identify the lymphoid regions proximal to the tumor, with BayesSpace providing higher resolution and more robust signal.

**Figure 3.**
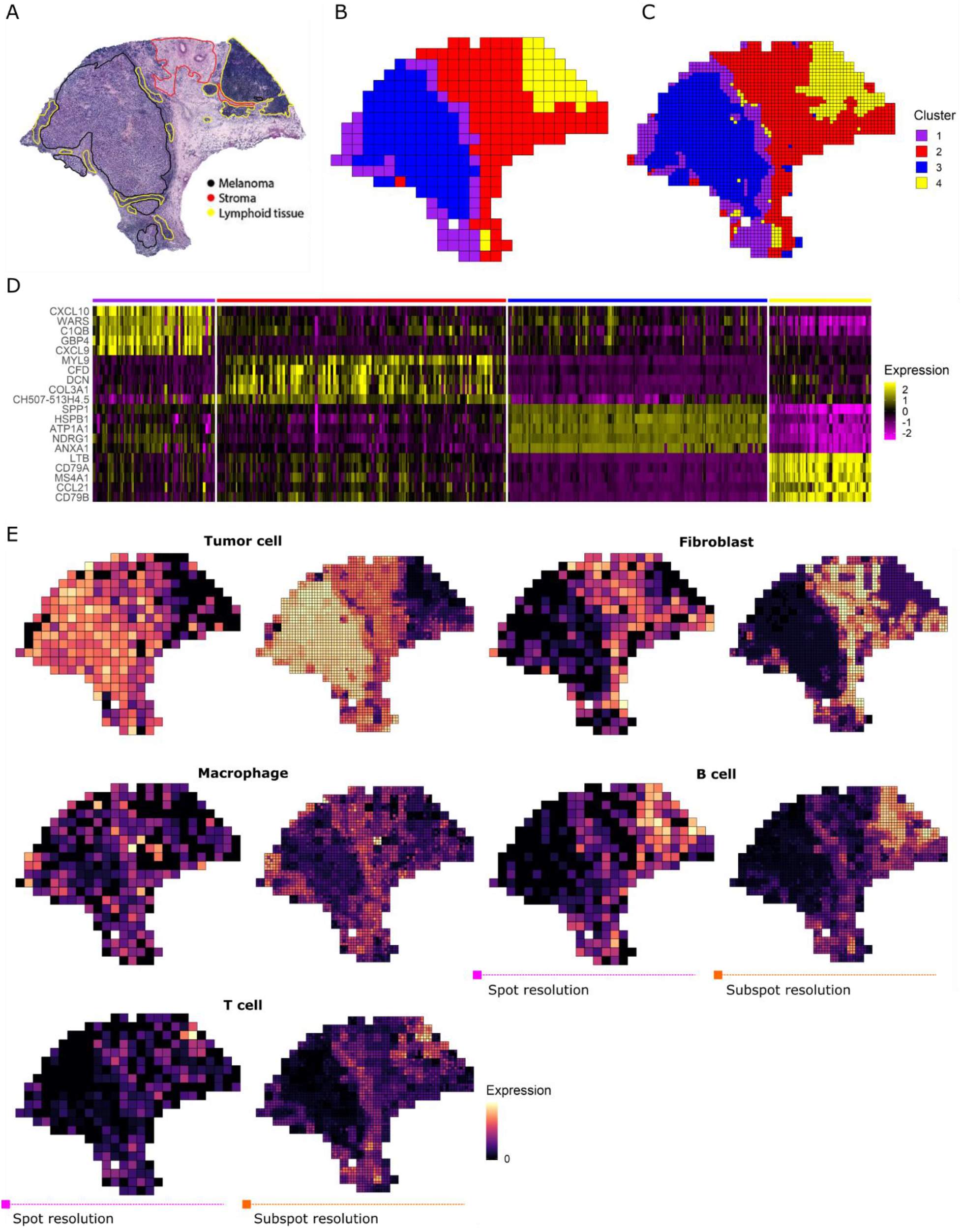
Enhanced resolution clustering identifies tumor-proximal lymphoid tissue in melanoma. (A) The original histopathological annotations of the H&E stained tissue find a section of melanoma (black) adjacent to tumor-proximal lymphoid tissue (yellow) and a region of stroma (red) separating these from a larger section of tumor-distal lymphoid tissue (yellow). Spatial clustering (B) and enhancement (C) generate biologically meaningful spatial domains corresponding to the original annotations. The enhanced resolution clustering identifies tumor-proximal lymphoid tissue (cluster 4; yellow) which is not resolved at spot-level clustering. (D) Differential expression analysis between the four clusters highlights the spatial differences in the expression of immune genes, cancer markers, and genes encoding extracellular matrix proteins. (E) For each of the five major cell types, the observed total spot-level expression (as measured by the summed log-normalized counts) of the defined marker genes (left) is shown alongside the corresponding enhanced resolution expression (right). We show the spatial expression plots for tumor cells (*PMEL*), fibroblasts (*COL1A1*), macrophages (*CD14*, *FCGR1A*, *FCGR1B*), B cells (*CD19*, *MS4A1*), and T cells (*CD2*, *CD3D*, *CD3E*, *CD3G*, *CD7*).

Using the enhanced PCs, we can generate high resolution maps of individual genes or expression profiles for major cell types as described in the Methods. Differential expression analysis performed on enhanced resolution gene expression indicates that the lymphoid regions have a distinct expression profile. We see elevated expression of lymphocyte markers such as *CD52* and *MS4A1* and lower expression of melanoma markers such as *MCAM* and *SPP1* relative to the surrounding tumor border (Fig. S2B). Enhanced resolution differential expression analysis between the four clusters highlights additional spatial variation in gene expression (Figure 3D). In the stroma (cluster 2), expression is higher for extracellular matrix proteins such as *DCN* and *COL3A1*. Furthermore, we reveal intra-tumor heterogeneity between the border and center of the tumor (clusters 1 and 3 respectively), with higher chemokine (*CXCL9*, *CXCL10*) activity at the border and elevated expression of genes related to cell proliferation (*HSPB1*) and metastasis (*ATP1A1*) at the center^22,23^.

We define tumor cell (*PMEL*), fibroblast (*COL1A1*), B cell (*CD19*, *MS4A1*), T cell (*CD2*, *CD3D*, *CD3E*, *CD3G*, *CD7*), and macrophage (*CD14*, *FCGR1A*, *FCGR1B*) expression profiles based on one or more marker genes from existing literature^24^. The enhanced expression profiles provide noticeably higher spatial resolution (Figure 3E). In particular, we can more clearly see immune expression on the periphery of the tumor. The contrast between *PMEL* expression in the tumor, stroma, and lymphoid tissue is also more apparent with enhanced resolution.

We also use BayesSpace to analyze a squamous cell carcinoma (SCC) Visium sample first described by Ji et al.^25^. A pathologist annotated the original H&E stained tissue to identify the tumor borders and other major tissue structures (Figure 4A). We defined expression profiles for the major cell types present in the sample based on known marker genes from literature: keratinocytes (*KRT1*, *KRT5*, *KRT10*, *KRT14*), melanocytes (*MLANA*, *DCT*, *PMEL*), myeloid cells (*LYZ*), and T cells (*CD2*, *CD3D*, *CD3E*, *CD3G*, *CD7*)^24,25^. The keratinocytes were further separated into basal keratinocytes (*KRT5*, *KRT14*) and suprabasal keratinocytes (*KRT1*, *KRT10*) since *KRT5* and *KRT14* form heterodimers that localize to the basal layer of the epidermis while *KRT1* and *KRT10* form heterodimers that localize to the suprabasal layer^26^. The enhanced spatial gene expression plots delineate the border between the basal and suprabasal layers noticeably more precisely than the spot-level plots (Figure 4B, Figure S3A).

**Figure 4.**
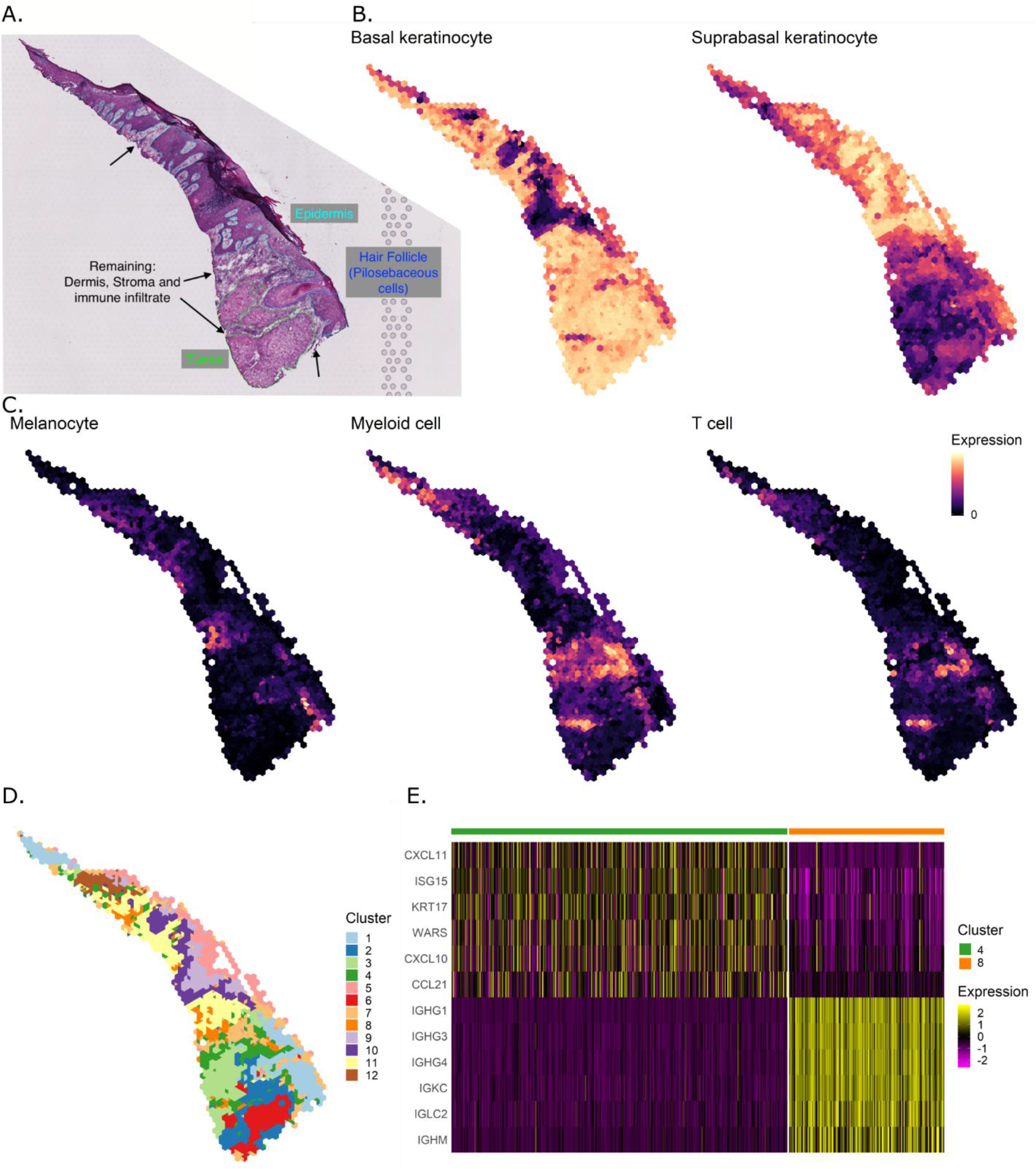
BayesSpace identifies transcriptionally distinct immune clusters. (A) Manual histopathological annotations of the stained H&E tissue differentiate the tumor (green), the epidermis (aqua), and a hair follicle (blue) from the remaining regions, which contain the dermis, stroma, and immune infiltrate. (B) Keratinocyte expression is high throughout the slide, but we see spatially distinct expression of basal (*KRT5*, *KRT14*) and suprabasal (*KRT1*, *KRT10*) keratinocytes at enhanced resolution. (C) Enhanced resolution maps of melanocyte (*MLANA*, *DCT*, *PMEL*), myeloid cell (*LYZ*), and T cell (*CD2*, *CD3D*, *CD3E*, *CD3G*, *CD7*) marker expression are biologically supported by the histopathological annotations. (D) The spots can be partitioned into twelve clusters, most of which display clear spatial patterns. (E) Clusters 4 and 8 have enriched immune cell expression but display substantial differences in expression of immunoglobulin genes and genes regulated by interferons.

The enhanced gene expression plots for melanocytes, myeloid cells, and T cells also better match the expected expression from the tissue structures as annotated (Figure 4C). Specifically, melanocytes are more clearly localized to the base of the hair follicle. Furthermore, the myeloid and T cells are more concentrated in the area annotated as stroma and immune infiltrate (Fig. S3A).

The high-resolution gene expressions are inferred from the enhanced clustering output of BayesSpace with twelve clusters (Fig. 4D). This number is chosen based on the elbow of the pseudo-log-likelihood by cluster number plot as a way to measure how well the clustering partition fits the data (Fig. S3B). We see high immune cell expression in clusters 4 and 8. Comparing these two clusters, differential expression analysis on the enhanced expression reveals that immunoglobulin genes are upregulated in cluster 8 (Fig. 4E). The marker genes for cluster 4, such as *CXCL10*, *CXCL11*, and *ISG15*, are induced by interferon-γ. This suggests that plasma cells are enriched in cluster 8 while inflamed epithelial cells are present in cluster 4, which is a region in close proximity to T cell markers that are likely driving this inflammation. We also compare clusters 2, 3, and 6, in which the tumor is located (Fig. S3C). In cluster 2, we see upregulation of *IGFBP2*, *IGFBP3*, and *IGFBP6*. These genes are members of the insulin-like growth factor-binding protein (IGFBP) family, which bind and stabilize *IGF1*, an important growth factor that can promote tumor growth^27^. *IGFBP3* is also a known marker of apoptosis^28^. Similarly to cluster 4, in cluster 3 the chemokine *CXCL10* is upregulated, a chemokine that has been shown to be associated with response to radiotherapy and survival in SCC patients^29^. Clusters 3 and 6 also show spatial expression of *LCE3D*, *SPRR2A*, *SPRR2D*, and *SPRR3*. These genes are members of the epidermal differential complex, which contains genes responsible for keratinocyte development and are upregulated in cutaneous SCC^30^.

### BayesSpace enhances gene expression patterns to near single-cell resolution on in silico spatial data

We conducted several simulations to demonstrate that BayesSpace clustering and enhancement outperform existing methods. In the first simulation where we simulated data modeled on our two experimental datasets (See Methods for details), the results show that BayesSpace spot-level clustering consistently outperforms all other methods in both the simulated melanoma and SCC datasets (Fig. 5A). *k*-means and Giotto clustering performed well in the melanoma sample and relatively poorly in the SCC sample. On the other hand, mclust performs relatively well across the two datasets.

**Figure 5.**
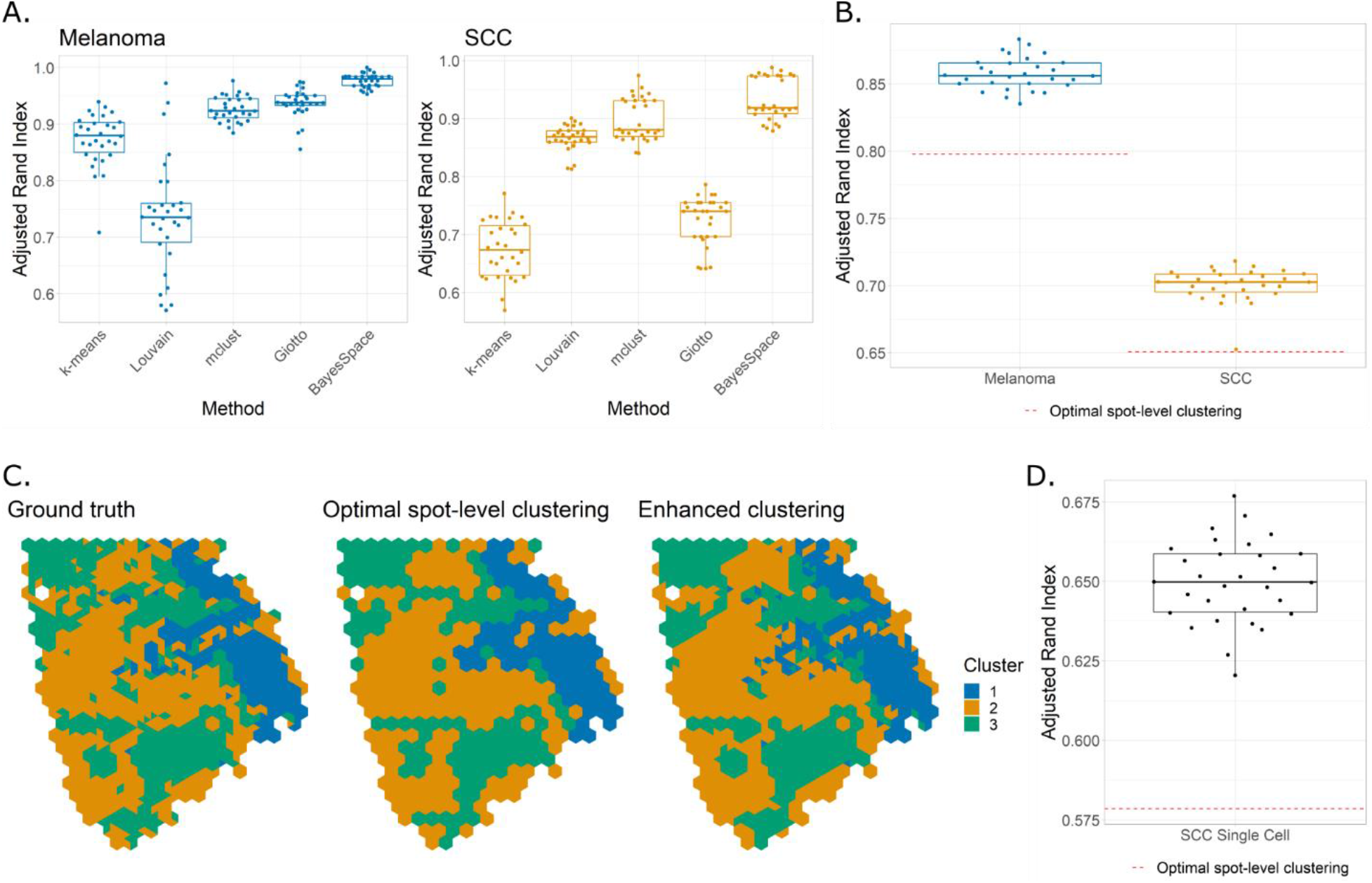
BayesSpace outperforms spatial and non-spatial clustering methods in simulated data. (A) In data simulated from the melanoma and SCC samples, BayesSpace spot-level clustering outperforms other clustering methods. (B) In simulations done at the sub-spot level, BayesSpace enhanced clustering outperforms the optimal spot-level clustering (red dotted line). (C) A single cell simulation is based on the ground truth from the enhanced SCC clusters (left). The optimal spot-level clustering for the ground truth can be compared to the output given by enhanced clustering. (D) The latter better approximates the ground truth, consistently across replicates in the simulation.

In the second simulation, we show that BayesSpace enhanced clustering outperforms the optimal clustering that can be achieved at the spot-level in melanoma and SCC samples that are simulated at the subspot level (Fig. 5B). In each dataset, the enhanced clustering ARI exceeds the optimal spot-level clustering in all thirty simulated replicates. This indicates that BayesSpace is able to increase the resolution of the data to better recapture finer details of the ground truth.

In the third simulation, we demonstrate that BayesSpace enhanced clustering can increase the resolution of data that are simulated from real, aggregated single cells (See Methods for details). BayesSpace better captures the spatial distribution of clusters than the optimal spot-level clustering, as illustrated in the spatial representation of the enhanced clustering results from one replicate (Fig. 5C). In regions with high mixing of clusters, there is little to no information available to resolve the cluster labels at the subspot level, but we see that BayesSpace is still often able to closely approximate the proportions of each cluster at the spot level. This shows that BayesSpace is able to detect isolated cells that would otherwise be missed due to the signal being diluted out from the aggregation of multiple cells at the spot level. The results further support our melanoma analyses where our enhanced analysis recovers lymphoid structure near the tumor that was not apparent at the spot level. In all, BayesSpace enhanced clusters better recapture the ground truth than the optimal spot-level clustering in all 30 replicates, again highlighting the consistency of our method (Fig. 5D) and showing that BayesSpace is able to enhance the spot level data to near single-cell resolution.

## Discussion

BayesSpace seamlessly integrates into the spatial transcriptomics analysis workflow by taking as input preprocessed data via the widely used SingleCellExperiment data structure. The output is likewise stored in a SingleCellExperiment object that can be used for downstream analyses. The methods are all conveniently implemented as an R package that has been submitted to Bioconductor. In all, BayesSpace provides an easy-to-use set of tools that facilitates the discovery of biological insights from spatial transcriptomics data.

We have demonstrated the utility of BayesSpace in identifying spatial clusters with similar expression profiles and enhancing the resolution of spatial transcriptomics. BayesSpace overcomes both the challenge in efficiently leveraging spatial information to inform the clustering of expression data and the limited resolution of current spatial transcriptomics technology. To our knowledge, BayesSpace is the first spatial transcriptomics model-based clustering method that uses a *t*-distributed error model to identify spatial clusters that are more robust to the presence of outliers caused by technical noise. Furthermore, our novel resolution enhancement method is the first to resolve spatial transcriptomics data at the subspot level without requiring the use of additional information besides spatial coordinates. This allows downstream differential expression analyses to compare finer and more biologically meaningful clusters. Importantly, the resolution enhancement approaches single cell resolution without the need for external single cell data. However, there is potential for the enhanced data to be integrated with external single cell data through deconvolution or label transfer methods. This represents a future direction of our research.

While our work focused on the ST and Visium platforms from 10X Genomics, BayesSpace should be applicable to other platforms where spots are arranged on a lattice. Slight modifications may be needed so that our spatial model can be used with a different neighborhood structure. Since BayesSpace models a lower dimensional representation of the data (i.e. PCA), it should also be applicable to other dimensional reduction techniques such as UMAP and possibly applied to other data types such as protein markers and multiomics. Finally, it may also be possible to extend BayesSpace to jointly cluster spots from multiple samples given appropriate data normalizations.

## Methods

### Data description

We applied BayesSpace to samples from three previously published spatial gene expression datasets, of which two are on the newer Visium platform. Details on dataset processing and availability are provided in the supplementary material. The first dataset included twelve human DLPFC samples from three individuals run on the Visium platform^4^. Briefly, each sample contained approximately 4,000 spots that were manually annotated to belong to one of the six DLPFC layers or the white matter. The second dataset involved melanoma samples run on the ST platform^2^. From this dataset, we analyzed the second replicate from biopsy 1 since it contained regions annotated as lymphoid tissue and was also described extensively in the original paper. Biopsy 1 contains 293 spots covered by tissue. The final dataset included ten human skin SCCs profiled on either the ST or the Visium platform^25^. Among the two samples run on the Visium platform, we chose to analyze patient 4 (P4) since the data quality was higher as shown in the original paper. Sample P4 contains 722 spots covered by tissue.

### Preprocessing and dimension reduction

In all datasets, raw gene expression counts were log-transformed and normalized using library size^31,32^. Principal component analysis (PCA) was then performed on the top 2,000 most highly variable genes. In downstream analyses, we modeled the top 15 principal components (PCs) from the DLPFC and SCC samples, and we modeled the top 7 PCs from the melanoma sample. In general, we recommend modeling the top 15 PCs to capture as much of the variability in the data as possible while limiting the rapid increase in space that occurs with higher dimensions. In the melanoma sample, many of the higher PCs exhibited higher numbers of extreme outliers (Fig. S5), with significant less variance suggesting that they most likely represent technical variability. Since the older ST technology has lower coverage and sequencing depth, fewer PCs are necessary for modeling.

### Spatial clustering model

BayesSpace implements a fully Bayesian model with a Markov random field (MRF) prior to encourage spots of the same cluster to be close to one another. Such models have been widely used in image analysis, including analyses of microarray images^15,16^. ST and Visium spots are arranged on square and hexagonal lattices, which provide a natural way to define a neighborhood structure (Fig. 1B). For each spot *i*, a low *d*-dimensional representation ***y***_*i*_ (e.g. PCs) of the gene expression vector can be obtained. We model the data as follows,

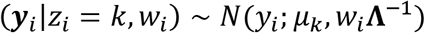

*z*_*i*_ ∈ {1, …, *q*} denotes the latent cluster that *i* belongs to, ***μ***_*k*_ and **Λ** denote the mean vector and precision matrix for cluster *k* and *w*_*i*_ is an unknown (observation-specific) scaling factor. The number of clusters *q* is determined by prior biological knowledge when available or otherwise by the elbow of the pseudo-log-likelihood plot (Fig. S3B). We place the following priors on ***μ***_*k*_, **Λ**, and *w*_*i*_:

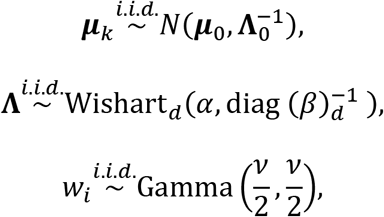

where ***μ***_0_, **Λ**_0_, *α* and *β* are fixed hyperparameters. By default, we set ***μ***_0_ to be the empirical mean vector of the data, which is the generally the zero vector for PCA input. **Λ**_0_ is set to 0.01 times the identity matrix to provide a weak prior that will be dominated by the data when there are spots assigned to the cluster. Similarly, we set *α* = 1 and *β* = 0.01 to provide a weak prior for the precision matrix. *v* denotes a fixed degrees of freedom parameter to control the heaviness of tails and is set to *v* = 4, which has been previously been shown to overcome the influence of outlier spots during clustering^16^. We also assume ***y***_***i***_ and *w*_*i*_ are independent, as such when marginalizing over *w*_*i*_ our normal likelihood becomes a multivariable *t* distribution with 0 mean and covariance matrix 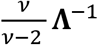. This formulation allows us to use a simple Gibbs sampling for updating most of the parameters, since the observations are normally distributed when conditioning on *w*_*i*_. The *w*_*i*_’s can also be interpreted as weights; the model will simply estimate a small weight value for any potential outlying data value. This provides robustness against outliers that can be commonly encountered in these types of data. Estimation of the parameters is done using a Markov chain Monte Carlo (MCMC) method. We initialize ***z*** using a non-spatial clustering method such as mclust by default^20^. Alternative initializations can also be supplied as a label vector. Then, iteratively and sequentially, each ***μ***_*k*_, **Λ**, and *w*_*i*_ is updated via Gibbs sampling and each *z*_*i*_ is updated via Metropolis-Hastings as described in supplementary material.

### Spatial clustering model at enhanced resolution

Relative to the clustering method, the model specification and parameter estimation is largely similar for the enhanced resolution clustering, though the unit of analysis is now a subspot rather than a spot. We segmented each ST spot into 9 subspots, while each Visium spot is segmented into 6 subspots (Fig. 1B). For ST, we used 9 subspots to help increase the resolution of the data, which is much lower than that of the Visium platform. Since gene expression is not observed at the subspot level, it is modeled as another latent variable that is also estimated through MCMC. The latent expression of each subspot *j* that is part of spot *i* is denoted 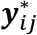, initialized to be ***y***_*i*_, and then updated via Metropolis-Hastings. In each iteration and for each spot, the new proposal is given by 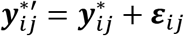 for each subspot such that ***ε***_*ij*_~*N*(**0**, ***σ***^2^*I*_*d*_) where *σ*^2^ is a small fixed parameter and ∑_*j*_ ***ε***_*ij*_ = **0**. In effect, this jitters the latent expression value of each subspot within a spot while keeping the total expression of the spot fixed. The proposal is accepted or rejected based on the conditional likelihood of the proposal given the other parameters. A weak Gaussian prior is placed on the latent expression to ensure that the jittered values do not drift too far away from ***y***_*i*_. Aside from replacing ***y***_*i*_ with 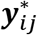, all other steps of the MCMC algorithm remain the same as in the spot-level clustering method.

### Mapping high-resolution PCs to high-resolution gene expression space

While BayesSpace can provide higher-resolution maps of spatial transcriptomic patterns, the modeling is done on the PC space and an additional step is necessary to map the principal component values back to the original log-normalized gene expression space. BayesSpace implements non-linear regression using XGBoost for predicting high-resolution gene expression^33,34^. A model is trained for each gene where the outcome is the measured gene expression at the spot level, and the predictors are the PCs generated from the original data. The fitted model can then be used to predict gene expression from the high-resolution PCs estimated using enhanced resolution clustering. The enhanced gene expression values can be visualized spatially and analyzed via differential expression methods (Fig. 1A). In our analyses, we use the Wilcoxon rank-sum test as implemented in Seurat to identify the top differentially expressed genes and also use Seurat for heatmap visualizations of the centered and scaled gene expression values^35^.

### Simulations

Using several simulations, we evaluate the performance of BayesSpace. The first simulation compares BayesSpace spot-level clustering to other non-spatial and spatial clustering methods: k-means, Louvain, mclust, and Giotto. We could not evaluate stLearn in simulation due to the need for an image as input. The simulated data are based on the melanoma and SCC samples introduced in the earlier results. Thirty replicates of simulated melanoma and SCC PCs are generated from *t*-distributions with means, precision, and spot labels determined by the spot-level clustering results of the real melanoma and SCC samples respectively (Fig. 3B, Fig. S4A). Other clustering methods are implemented as described in the supplement with the true cluster number provided as input when possible. Performance is assessed using the adjusted Rand index between the ground truth spot labels and the clustering results.

In the second simulation, we evaluate the performance of BayesSpace subspot-level enhanced clustering. As before, we simulate 30 replicates from *t*-distributions with means, precision, and labels based on the real melanoma and SCC samples, but in this simulation we generate subspots using the enhanced clustering results as the ground truth (Fig. 3C, Fig. 4D). The simulated subspot-level PCs are averaged to provide spot-level PCs that are given as input to BayesSpace. We can use the modal ground truth label of the subspots within each spot to generate an optimal spot-level clustering for each dataset (Fig. S4B). The ARI between this optimal spot-level clustering and the subspot-level ground truth represents the highest ARI that can be achieved when all subspots within a spot must belong to the same cluster, as is the case with spot-level clustering.

In the third simulation, we sample data from real single cells rather than simulating PCs. Ji et al. provide matching scRNA-seq data for the SCC sample^25^. These single cells can be sampled into subspots, providing another way to evaluate the performance of enhanced clustering without relying on model-based data generation. Given the limited number of single cells and distinct cell types, we use only the positions from the lower portion of the SCC sample and collapse the original clusters into three (Fig. 5C). Clusters 1, 2, and 3 are sampled from melanocytes, myeloid cells, and epithelial cells respectively in the scRNAseq data. The subspots are aggregated into spots by averaging the log-normalized count of each gene, and then processed to generate PCs as described for real data in the Methods. The optimal spot-level clustering is again inferred from the modal cluster label of subspots within each spot (Fig. 5C) and is used to initialize the BayesSpace enhanced clustering.

## Code availability

The BayesSpace software package has been submitted to Bioconductor, and the source code is publicly available at https://github.com/edward130603/BayesSpace.

## Data availability

The datasets analyzed in this paper are available in raw form through their original authors (see details in Supplementary Information), and the SingleCellExperiment objects we prepared for analysis with BayesSpace are available through the BayesSpace package.

## Acknowledgements

This research was supported by funding from National Institutes of Health U19AI128914 and P01CA225517 to R.G. and P.N. and the Immunotherapy and Data Science Integrated Research Centers at Fred Hutch to E.Z., M.S., X.R. and J.H.B.

We thank Dr. Ata Moshiri from the UW Division of Dermatology for his review of T.P.’s histopathological annotations; and Dr. Quan Nguyen and Xiao Tan at the University of Queensland for their assistance in applying stLearn.

## Author contributions

E.Z. and R.G. formulated the method and wrote the manuscript. M.S. and E.Z. developed the software. E.Z., M.S., and X.R. analyzed the data. T.P. contributed to annotation and interpretation of cancer samples. P.N., J.B., and R.G. supervised the project.

