## Supplementary Figures for "BayesSpace enables the robust characterization of spatial gene expression architecture in tissue sections at increased resolution"

**Figure S1.** The spatial distribution of $w_i$'s are shown for sample 151673. Notice that many of the visible outlier spots seen in PC2 and PC3 (some of which are annotated by the green oval and green arrows) are downweighted by BayesSpace as indicated by the $w_i$ value. The corresponding locations on the $w_i$ plot are annotated in red.

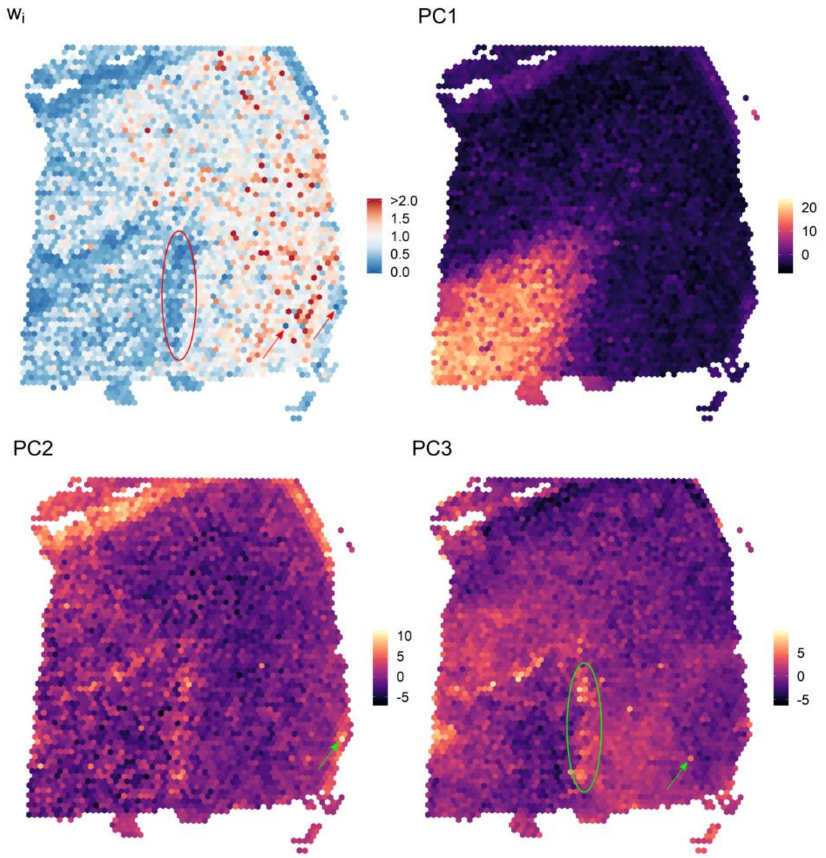

**Figure S2.** (A) The results of other clustering methods are shown. Using default parameters, Louvain clustering partitions the spots into five clusters. For the other methods, the spots are partitioned into four clusters to match the BayesSpace results in Figure 3. (B) The expression of lymphoid regions identified near the tumor are compared to the expression of the remaining tumor border (left). The analysis reveals that lymphocyte markers are expressed more in the areas identified as lymphoid tissue (right).

A.

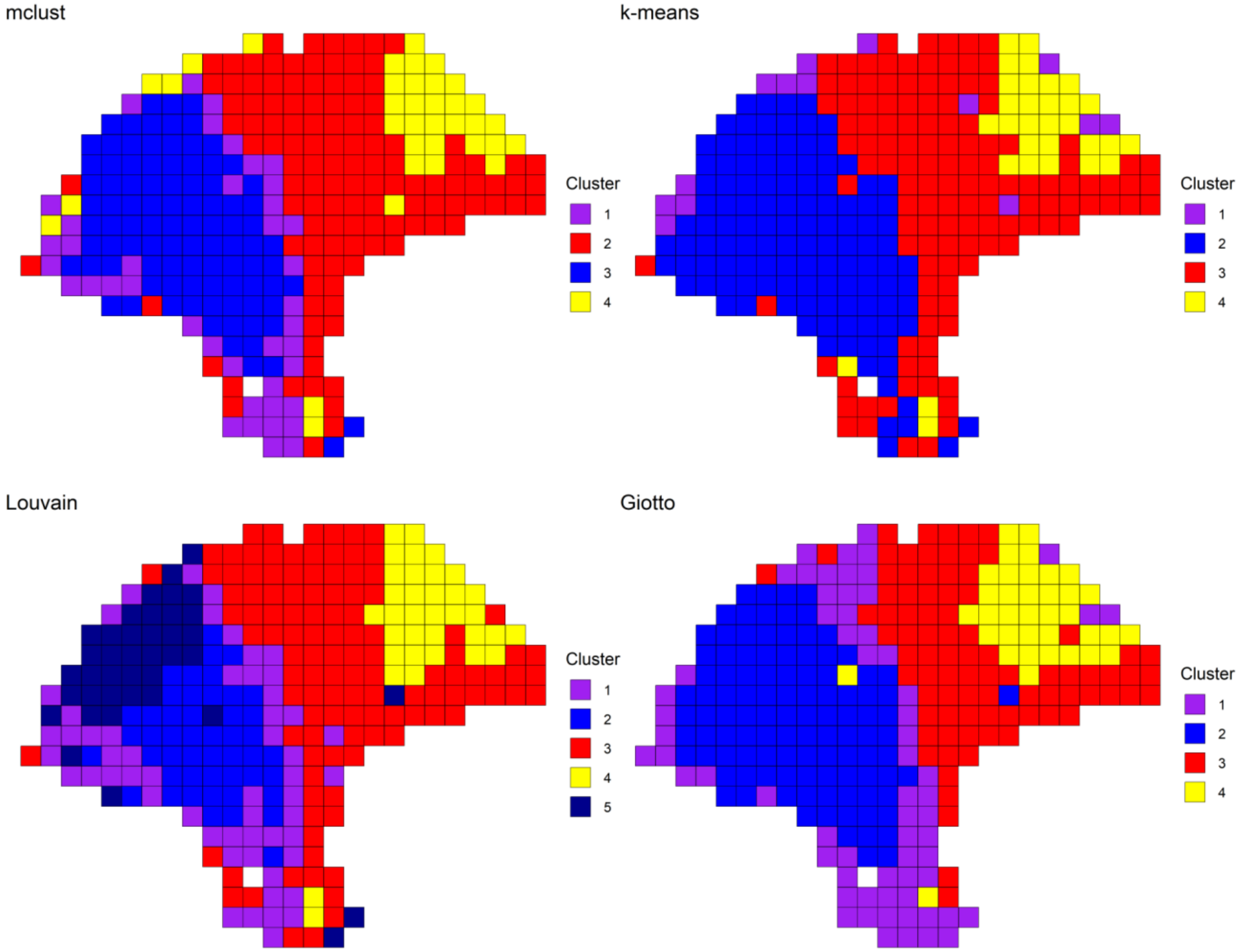
mclust
Cluster
1
2
3
4
k-means
Cluster
1
2
3
4
Louvain
Cluster
1
2
3
4
5
Giotto
Cluster
1
2
3
4

B.

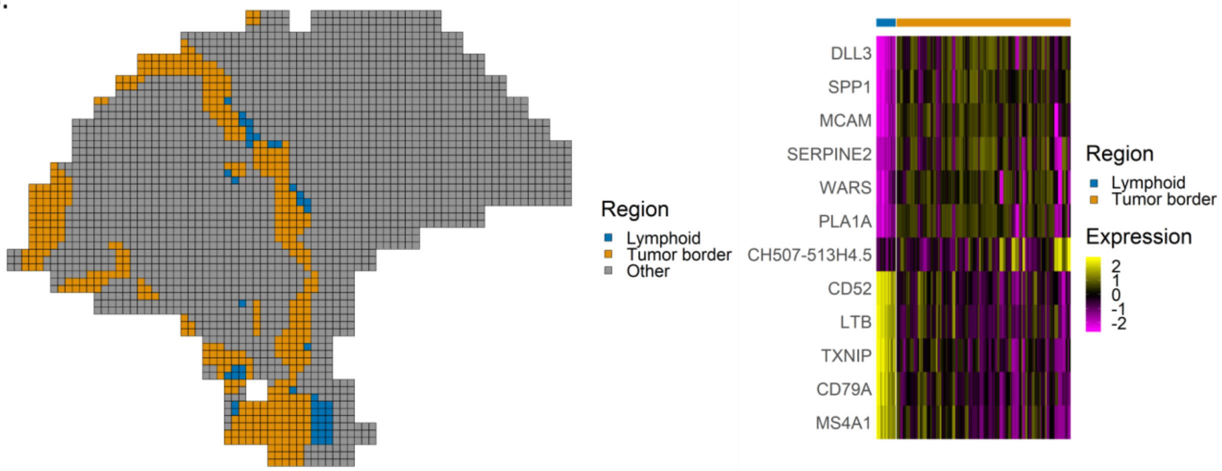
Region
Lymphoid
Tumor border
Other
DLL3
SPP1
MCAM
SERPINE2
WARS
PLA1A
CH507-513H4.5
CD52
LTB
TXNIP
CD79A
MS4A1
Region
Lymphoid
Tumor border
Expression
2
1
0
-1
-2

**Figure S3.** (A) The spot-level marker expression of basal keratinocytes, melanocytes, and myeloid cells are shown. The areas in red boxes are also shown at enhanced resolution in the inset within the black box. In each case, enhanced resolution spatially refines the marker expression. (B) `spatialCluster()` is run for 1,000 iterations with number of clusters set between 3 and 20. The mean negative pseudo-log-likelihood over iterations (excluding a 100 iteration burn-in) is plotted for each cluster number. $q = 12$, denoted by the vertical red line, is a reasonable choice for the elbow. (C) Differential expression analysis between the three tumor clusters highlights heterogeneity within the tumor.

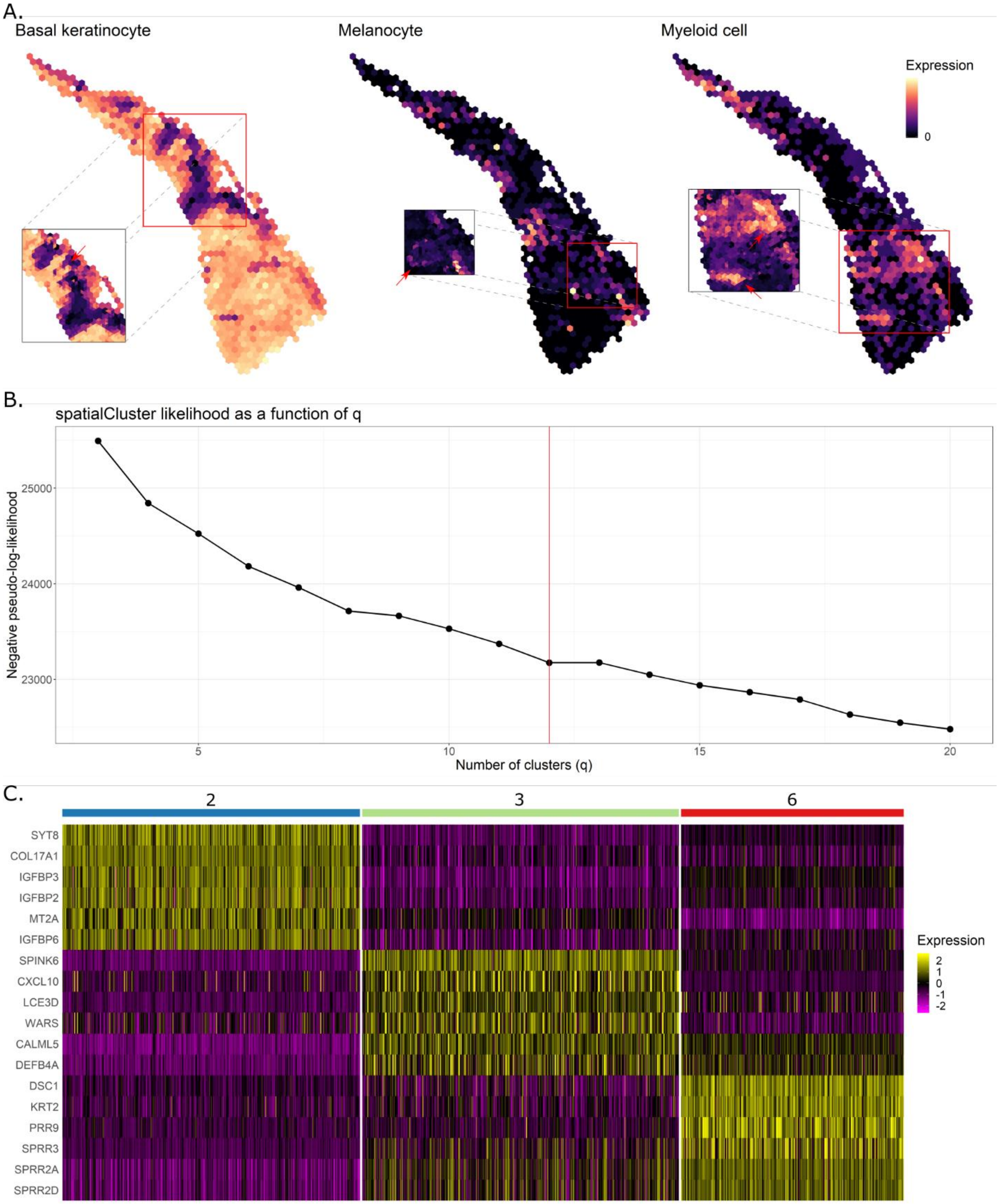

A.
Basal keratinocyte
Melanocyte
Myeloid cell
Expression
0
B.
spatialCluster likelihood as a function of q
Negative pseudo-log-likelihood
25000
24000
23000
5
10
15
20
Number of clusters (q)
C.
2
3
6
SYT8
COL17A1
IGFBP3
IGFBP2
MT2A
IGFBP6
SPINK6
CXCL10
LCE3D
WARS
CALML5
DEFB4A
DSC1
KRT2
PRR9
SPRR3
SPRR2A
SPRR2D
Expression
2
1
0
-1
-2

**Figure S4.** (A) The spot-level BayesSpace clustering partition of the SCC sample is shown. (B) The optimal spot-level clustering partitions for the melanoma and SCC enhanced clustering simulations are shown.

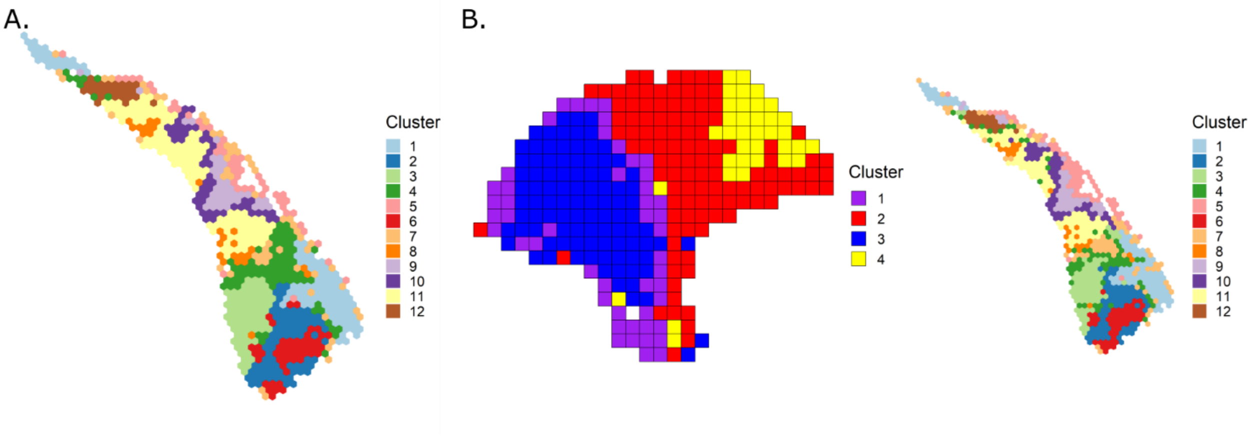

**Figure S5.** The higher PCs of the melanoma sample have large outliers. Only PCs 1 through 7 are used in modeling the melanoma sample to capture as much biological variation as possible while limiting the impact of outliers. For reference, the distribution of PCs from DLPFC sample 151673 and the SCC sample are shown below. These Visium samples do not have large outliers in the higher PCs.

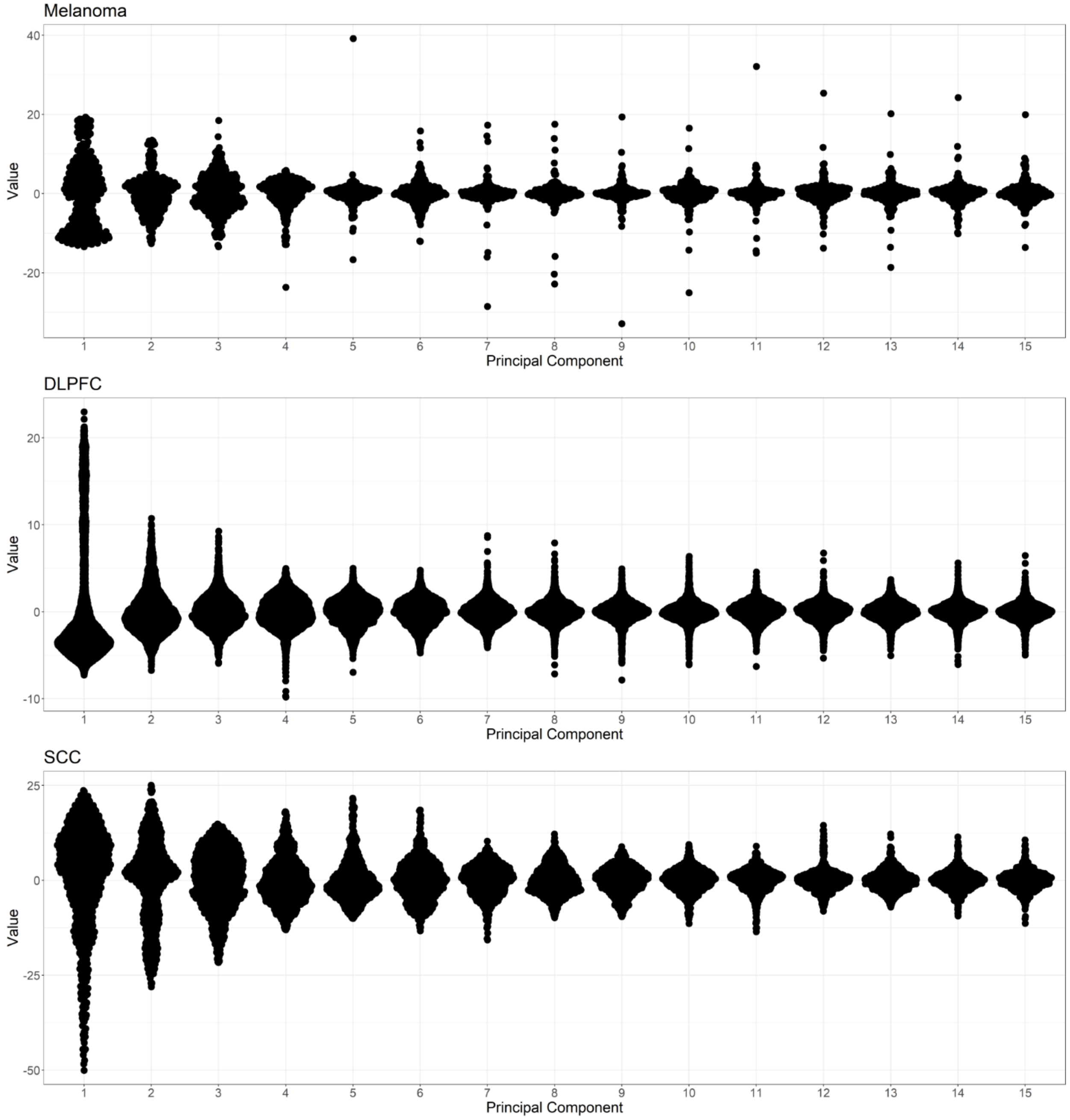
