## Supplementary Notes for "BayesSpace enables the robust characterization of spatial gene expression architecture in tissue sections at increased resolution"

*Dataset processing and availability*

- Dorsolateral prefrontal cortex[1]
  - Spatial data (Maynard et al. 2020). A SingleCellExperiment object containing the counts and spatial coordinates for all samples was downloaded using the `spatialLIBD::fetch_data()` method and then subset by sample.
- Melanoma[2]
  - Spatial data (Thrane et al. 2018). Counts matrices were downloaded from the Spatial Research website (https://www.spatialresearch.org/wp-content/uploads/2019/03/ST-Melanoma-Datasets_1.zip). Array row/column coordinates were obtained from the column labels of these matrices. An aligned image and pixel coordinates were not available for this sample.
- Squamous cell carcinoma[3]
  - Spatial data (Ji, Rubin, et al. 2020). A counts matrix containing expression profiles from all samples and tissue positions were downloaded from GEO (accession GSE144239).
  - Single-cell data (Ji, Rubin, et al. 2020). The counts matrix containing the matched single-cell experiment and a table of cell type annotations were downloaded from GEO (accession GSE144236).

After downloading each dataset, we performed basic data cleaning to convert each expression matrix into a SingleCellExperiment[4] object with array and pixel coordinates stored in the object's `colData`. PCA was performed on the log-normalized expression of the top 2,000 highly variable genes in each dataset.

Snakefiles[5] and scripts used to process each dataset are available at https://github.com/msto/spatial-datasets, and processed datasets are accessible via the `BayesSpace::getRDS()` method.

*Comparison of other clustering methods*

We applied the other clustering methods as follows:

- *k*-means: The base R `stats::kmeans()` function is used with default parameters.
- mclust: The "EEE" model is used to match the model used by BayesSpace. All other parameters are kept at the defaults.
- Louvain: The shared nearest neighbors graph is constructed via `scran::buildSNNGraph()` using Jaccard weights and the default 10 neighbors. Louvain clustering is done via `igraph::cluster_louvain()`
- Giotto/HMRF: We adapted the Giotto workflow described in their online tutorial (http://spatialgiotto.rc.fas.harvard.edu/giotto.visium.brain.html). Specifically, we filtered genes expressed in fewer than 10 spots after encountering numerical issues without this filter. We used Giotto's internal functions to normalize expression, identify HVGs and perform PCA, and create a Delaunay network. The Delaunay network was created with maximum distance set to 'auto', which reproduced our defined neighborhood structure. Finally we used `Giotto::doHMRF()` to obtain spatial domains (cluster assignments) from spatially expressed genes (using k-means binarization; `Giotto::binspect()`) in the DLPFC samples and the top 15 PCs in the remaining samples. The parameter beta, which controls the strength of interaction between spots, was set to 2 for the melanoma sample and simulations, 3 for the SCC sample and simulations, 9 for the DLPFC samples. The DLPFC sample beta parameter was higher due to the higher dimensionality

of the input (genes rather than PCs). We chose to use genes rather than PCs for this dataset due to the poor performance of Giotto on PCs here.

- stLearn: stLearn was applied to the DLPFC samples as described in their online tutorial (https://stlearn.readthedocs.io/en/latest/stSME_clustering.html#Human-Brain-dorsolateral-prefrontal-cortex-(DLPFC)). We applied stLearn once using the 63 marker genes referenced from the original study (https://github.com/LieberInstitute/HumanPilot/blob/master/mouse_layer_marker_info_cleaned.csv) and once using the 2,000 highly variable genes upon which we performed PCA.

*Formal description of the Markov chain Monte Carlo method*

We update $\boldsymbol{\mu}_k, \boldsymbol{\Lambda}_k$, and $w_i$ via Gibbs sampling from the posterior distributions:

$$\boldsymbol{\mu}_k \sim N\left(\left(\boldsymbol{\Lambda}_0 + \boldsymbol{\Lambda} \sum_{i:\{z_i=k\}} w_i\right)^{-1} \left(\boldsymbol{\Lambda}_0 \boldsymbol{\mu}_0 + \sum_{i:\{z_i=k\}} w_i \boldsymbol{y}_i\right), \left(\boldsymbol{\Lambda}_0 + \boldsymbol{\Lambda} \sum_{i:\{z_i=k\}} w_i\right)^{-1}\right)$$

$$\boldsymbol{\Lambda} \sim \text{Wishart}_d\left(\sum_i w_i + \alpha, \left(\text{diag}(\beta)_d + \sum_i w_i (\boldsymbol{y}_i - \boldsymbol{\mu}_i)^T (\boldsymbol{y}_i - \boldsymbol{\mu}_i)\right)^{-1}\right)$$

$$w_i \sim \text{Gamma}\left(\frac{d+\nu}{2}, \frac{\nu + (\boldsymbol{y}_i - \boldsymbol{\mu}_i)^T \boldsymbol{\Lambda} (\boldsymbol{y}_i - \boldsymbol{\mu}_i)}{2}\right)$$

Given $n$ spots and a set cluster number $q$, the cluster label vector $\boldsymbol{z}$ can take values in $\{1, \dots, q\}^n$ and the Metropolis-Hastings algorithm is used to explore this parameter space. For each spot $i$, a new cluster label $z_i'$ is proposed from $\{1, \dots, q\} \backslash z_i$ and accepted or rejected based on the ratio of posterior distributions $\frac{p(z_i|y_i)}{p(z_i'|y_i)} = \frac{p(y_i|z_i)\pi(z_i)}{p(y_i|z_i')\pi(z_i')}$, where the likelihood $p(y_i|z_i)$ is given by

$(\boldsymbol{y}_i|\boldsymbol{\mu}_k, \boldsymbol{\Lambda}, z_i = k, w_i) \sim N(\boldsymbol{\mu}_k, w_i^{-1}\boldsymbol{\Lambda}^{-1})$ and the spatial smoothing prior is given by the Potts model:

$$\pi(z_i) = \exp\left(\frac{\gamma}{|\langle i\ j\rangle|} \times 2 \sum_{\langle i\ j\rangle} I(z_i = z_j)\right),$$

Where $\langle i\ j\rangle$ denotes all spots $j$ that are neighbors of $i$, and $\gamma$ is a fixed parameter controlling the strength of the smoothing. Given that ST and Visium spots are arranged on a regular lattice, there is a natural way to define spatial neighbors (Fig. 1B). ST spots can have up to 4 neighbors while Visium spots can have up to 6. By default, we use $\gamma = 2$ for ST and $\gamma = 3$ for Visium, accounting for the higher number of neighbors.

Estimation of the parameters is done using a Markov chain Monte Carlo (MCMC) method. We initialize $\boldsymbol{z}$ using a non-spatial clustering method such as *k*-means or mclust. Then, iteratively and sequentially, each $\boldsymbol{\mu}_k, \boldsymbol{\Lambda}$, and $w_i$ is updated via Gibbs sampling and each $z_i$ is updated via Metropolis-Hastings as described above. After iterating for a fixed number of iterations, the mode of the chain (after discarding a specified number of burn-in iterations) for each $z_i$ is assigned as the cluster label for the corresponding spot $i$.

Note that $(\boldsymbol{y}_i|\boldsymbol{\mu}_k, \boldsymbol{\Lambda}_k, z_i = k) \sim T_\nu(\boldsymbol{\mu}_k, w_i^{-1}\boldsymbol{\Lambda}^{-1})$, a multivariate *t*-distribution with a fixed $\nu$ degrees of freedom which allows for a more robust error model. In the BayesSpace software package, we additionally provide a simplification of the model where all weights are fixed to be $w_i = 1$, resulting in Gaussian marginal errors.
